# Improving bioinformatics prediction of microRNA targets by ranks aggregation

**DOI:** 10.1101/224915

**Authors:** Aurélien Quillet, Chadi Saad, Gaētan Ferry, Youssef Anouar, Nicolas Vergne, Thierry Lecroq, Christophe Dubessy

## Abstract

microRNAs are non-coding RNAs which down-regulate a large number of target mRNAs and modulate cell activity. Despite continued progress, bioinformatics prediction of microRNA targets remains a challenge since available softwares still suffer from a lack of accuracy and sensitivity. Moreover, these tools show fairly inconsistent results from one another. Thus, in an attempt to circumvent these difficulties, we aggregated all human results of three important prediction algorithms (miRanda, PITA and SVmicrO) showing additional characteristics in order to rerank them into a single list. This database is freely available through a webtool called miRabel (http://bioinfo.univ-rouen.fr/mirabel/) which can take either a list of miRNAs, genes or signaling pathways as search inputs. Receiver Operating Characteristic curves and Precision-Recall curves analysis carried out using experimentally validated data and very large datasets show that miRabel significantly improves the prediction of miRNA targets compared to the three algorithms used separatly. Moreover, using the same analytical methods, miRabel shows significantly better predictions than other popular algorithms such as MBSTAR and miRWalk. Interestingly, a F-score analysis revealed that miRabel also significantly improves the relevance of the top results. The aggregation of results from different databases is therefore a powerful and generalizable approach to many other species to improve miRNA target predictions. Thus, miRabel is an efficient tool to accurately identify miRNA targets and integrate them into a biological context.

## Introduction

Mature microRNAs (miRNAs) are unpolyadenylated and uncapped 21-23 nucleotides endogenous non-coding single strand RNAs. They act at the post-transcriptional level to regulate gene expression in eukaryotic organisms. At least 60% of human genes are believed to be regulated by miRNAs as shown by a genome wide analysis [1]. Since their discovery in 1993 [2], it has been clearly established that miRNAs act as key regulators of several cell processes such as proliferation, differentiation, metabolism and apoptosis [3]; it is therefore not surprising to find them involved in numerous pathophysiological processes [4]. To date, 2,588 mature miRNAs (http://www.mirbase.org/) have been identified in human and each of them has the ability to potentially regulate several hundred of target mRNAs and each targeted mRNA can be regulated by tens of miRNAs [5], thus creating a large and complex regulation network of gene expression unsuspected before. They work mostly through imperfect base-pairing hybridization to mRNA, generally in the 3’-UTR [6], to block translation or rarely to induce mRNA degradation [7]. Moreover, it was shown that miRNA binding sites are also found in the 5’-UTR and in the coding region [8]. The bioinformatics identification of miRNA targets remains a challenge because mammalian miRNAs are characterized by a poor homology toward their target sequence except in the conserved “seed” region that mostly comprises nucleotides 2-7 of the miRNA [9]. Nevertheless, several algorithms have been developed in recent years in order to include a set of features known to modulate the interaction between miRNA and their cognate mRNA in addition to the essential Watson-Crick pairings [10]. Among them, the most relevant are the free energy of the miRNA::mRNA system [11], the conservation of sequences among species [12] and the accessibility of binding sites [13]. This resulted in the creation of more than 105 target prediction tools (as of November 2017, from OMICtools’ database [14]), all of which have their strengths and weaknesses [15, 16].These tools are useful to reduce the amount of potential targets in order to streamline the experimental validations [17]. However, their predictions suffer from a poor accuracy and sensitivity as revealed by experimental data [18, 19]. In addition, computational results are very divergent depending on how the bioinformatics tools take into account the aforementioned features of miRNA::mRNA interactions [20]. Moreover, several studies clearly show that algorithms performances depend on the dataset used [21, 22]. So far, no single method consistently outperforms others in the miRNA targets prediction field, thus supporting the idea that databases content combination is an efficient way to improve MTI prediction. Assuming that an interaction predicted by more than one algorithm is more likely to be functional, databases such as miRWalk [23, 24], miRSystem [25], miRGator [26] or, more recently, Tools4miRs [27], store and/or compare results predicted by several popular tools using statistics and mRNA/protein expression data. Ritchie et al. [28], however, demonstrated that targets resulting from the intersection of two lists of predictions are not more likely to be present in the intersection of two other lists. Therefore, intersecting results does not increase the probability of retaining true positives. Moreover, approaches based on intersection of predictions may lead to decreased sensitivity because of possibly omitting valid interactions as shown by Sethupathy et al. [29]. In order to circumvent these limitations, we proposed to compute a new unique score based on the aggregation of the interaction ranks taken from other well known prediction algorithms. To test our hypothesis, we aggregated three major prediction algorithm results which enabled us to show that this new score significantly improves miRNA targets prediction compared to other prediction tools. To allow a more comprehensive analysis, the results of this aggregation were eventually linked to their respective cellular pathways using KEGG database, and implemented in a web tool named miRabel. Interestingly, miRabel can take either a list of miRs, genes or pathways as search inputs and retrieve the linked results.

## Materials and methods

### Aggregated databases

Computationally predicted human miRNA::mRNA interaction databases generated by miRanda [30], PITA [31] and SVMicrO [32] were used. These publicly available online algorithms have been chosen because each of them uses different and complementary features of miRNA::mRNA interactions such as seed match, interspecies conservation, free energy, site accessibility and target-site abundance (Table S1) [10]. The ranks of each predicted interaction retrieved from one or more of these databases have been aggregated using the R package RobustRankAggreg (RRA) (v1.1) [33] with R (v3.2.0). The new score resulting from the aggregation is used to re-rank each interaction and also indicates the significativity of the proposed rank in miRabel.

### Testing datasets

Two types of testing datasets were used for each of the comparisons described in this paper. First, to compare the different aggregation methods, we used one million randomly selected interactions within aggregated data. Validated interactions accounted for 3% of the testing dataset. For the other evaluations, all common interactions between compared databases were used (Fig.1A). It resulted in extremely large datasets (>500,000 interactions) which reduced the amount of possible analysis due to computation time (several weeks). This led us to design a second type of datasets of 50,000 interactions randomly picked from the corresponding larger dataset. For each large dataset, 10 smaller ones were created (Fig.1B). The amount of experimentally validated interactions within these randomly picked ones was set so as to remain close in proportion to the main, larger dataset. These smaller datasets allowed us to increase the relevance and statistical significance of performance results.

**Figure 1:**
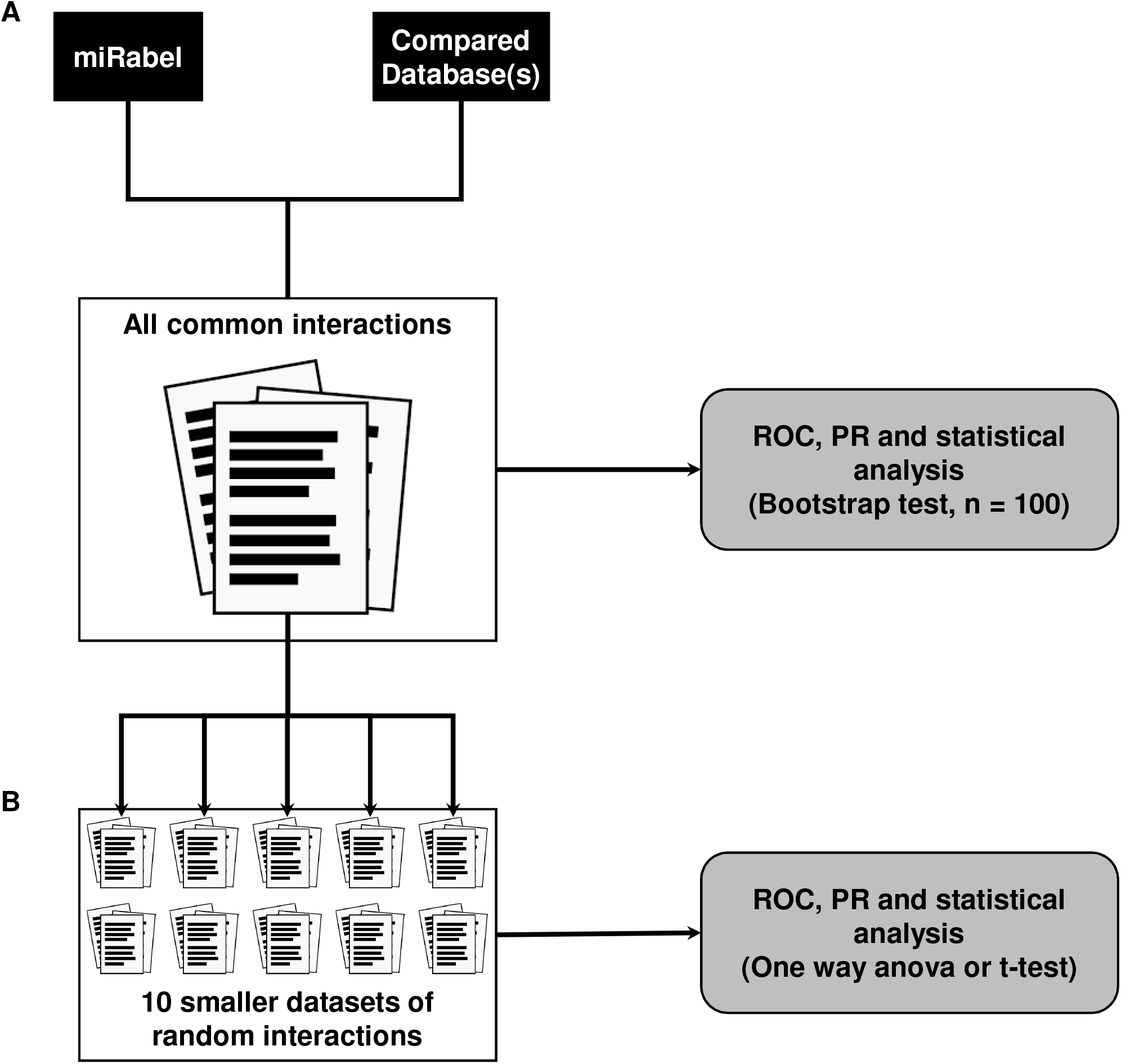
Testing datasets design and databases performance analysis methodology. A large dataset containing all common interactions between compared databases is created (**A**), For ease of use, 10 smaller datasets of 50,000 interactions were randomly picked from all common ones (**B**). Predictions performance are then compared using ROC and PR analysis on all datasets.

### Performance analysis methods

On each dataset, a receiver operating characteristic (ROC) analysis was done using the area under curve (ROC_AUC) as implemented in the R package pROC [34]. To analyse top prediction results, a specificity of 90% was set as a threshold in order to compute partial ROC (pROC90%) and the corresponding AUC (ROC_pAUC90%) and sensitivity. To focus on which classifier better identifies true positive interactions, datasets were further compared with precision and recall (PR) curves using R programming as well. For the same purpose as with the pAUC of the ROC analysis, we calculated the harmonic mean between the precision and the recall (F-score) for different percentages of the top interactions.

### Statistics

Statistical analysis of results obtained with smaller datasets were done using either a Repeated Measures One Way ANOVA with Dunnett’s post-test or a Student t-test depending on the number of compared groups with GraphPad Prism software (version 6.00 for Windows, GraphPad Software, La Jolla California USA).

## Results

### miRabel overview

#### miRabel: a database for microRNA target predictions

The database was designed with MySQL (http://www.mysql.com/) using InnoDB motor and includes predictions from miRanda [30], PITA (v.6.0) [31] and SVMicrO [32]. It contains tables for the 2,578 human miRNAs (for which 1,107 have target mRNAs), 20,532 genes and 275 pathways. This represents more than 8.6 million predicted interactions from which 123,373 are experimentally established. These experimentally validated interactions are taken from miRTarBase (v.6.0) [35] and miRecords [36], whereas 5’UTR and CDS predictions are retrieved from miRWalk database (v.2.0) [24]. Genes and pathways information as well as their relationships were retrieved from KEGG’s database while miRNA data were from miRBase (release 21, http://www.mirbase.org/) and linked with miRNA target predictions. Since the annotation of miRNAs has changed in the past few years, a conversion tool was developed to automatically convert the names of miRNA queries in the latest version used by miRBase. This tool is also accessible from the home page. In order to standardize gene names from the different tools, they were converted to the NCBI gene ID and a table containing their synonyms has been built. Potential interactions between miRNAs and genes were obtained based on our prediction method represented as shown in Fig. 2A. Pathways linked to the resulting interactions can be retrieved and ranked according to the proportion of its interactions regulated by a given miRNA. The number of validated interactions for this miRNA present in each pathway is also indicated.

**Figure 2:**
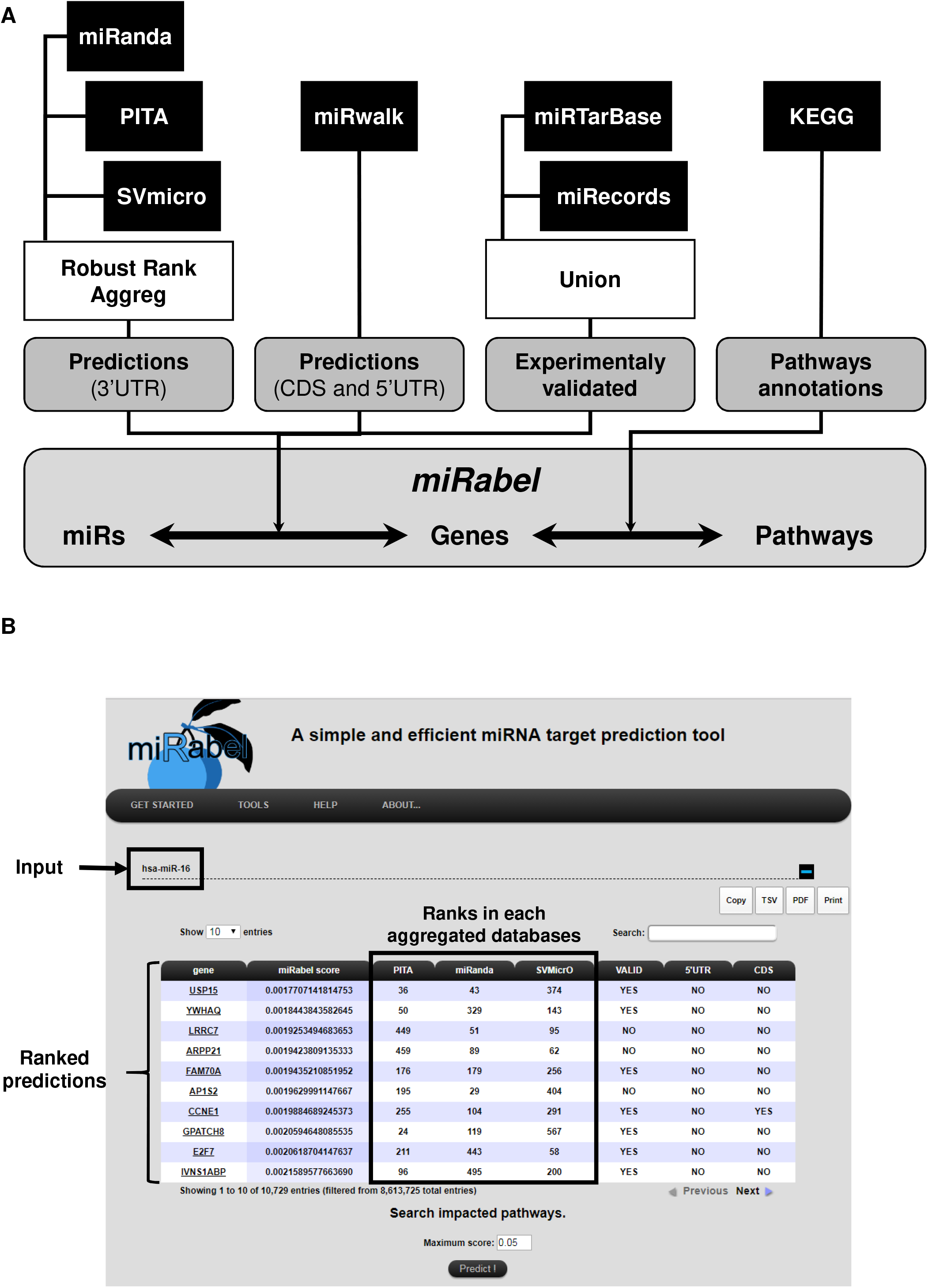
Overview of miRabel. Predictions results from miRanda, PITA and SVMicrO for 3’UTR are aggregated using Robust Rank Aggreg. 5’UTR and CDS predictions are retrieved from miRWalk database. Experimentally validated interactions are identified using miRTarBase and miRecords. Links between predictions and pathways are established based on KEGG information (**A**). An example of miRabel web interface is shown using predictions for hsa-miR-16. Predicted targets are ranked according to miRabel’s score. Rank found for this interaction in each database are indicated as well as its experimental validation status and mRNA sub-localization (**B**).

#### The web interface

The web interface was designed with PHP (http://www.php.net) and CSS (http://www.cssflow.com/). It enables users to query the system directly by miRNA name, by gene name or by pathway name (Fig. 2B). Multiple queries are allowed in order to identify common miRNAs, genes or pathways among the results. Alternatively, miRabel can be queried by uploading a text file containing the same information. Queries by pathways are easily made thanks to asynchronous database queries and name completion. The results are visualized by using the DataTable plugin of the JQuery framework which allows to create tables that can be easily filtered and sorted. Genes are linked to their NCBI gene homepage using their unique gene ID. Results can be copied, printed or exported in tabulated-separated or pdf formats. An online documentation section is also provided to help users in their searches. MiRabel website can be found at http://bioinfo.univ-rouen.fr/mirabel/.

### Evaluating aggregation methods

The performances of the aggregation methods (Mean, Default (i.e. RobustRankAggreg, RRA), Geometric mean, Median, Min, Stuart) provided by the R package RRA have been compared to each other (except for the Stuart method due to extensive computation time). ROC and PR analysis show that the mean of the ranks provides the best result (ROC_AUC_Mean_ = 0.5790, PR_AUC_Mean_ = 0.0436) (Fig. 3A-D). Interestingly, the F-score for different percentage of the top interactions indicates that the mean method is also the most consistent in promoting validated interactions (Fig. 3E-F). These results were confirmed using 10 smaller datasets. There again, the mean of the ranks provides the best results (ROC_AUC_Mean_ = 0.6888±0.0030, PR_AUC = 0.0290±0.0006) with significant statistical differences compared to other proposed methods (Table S2). When looking at top predictions only, the mean method remains significantly better than other compared methods (Table S1). Moreover these analyses show that among the ten datasets, the mean aggregation method provides the best ROC_AUC nine times whereas geometric mean method succeeds only one time (data not shown). These results led us to use the mean method to aggregate the ranks of miRanda, PITA and SVMicrO.

**Figure 3:**
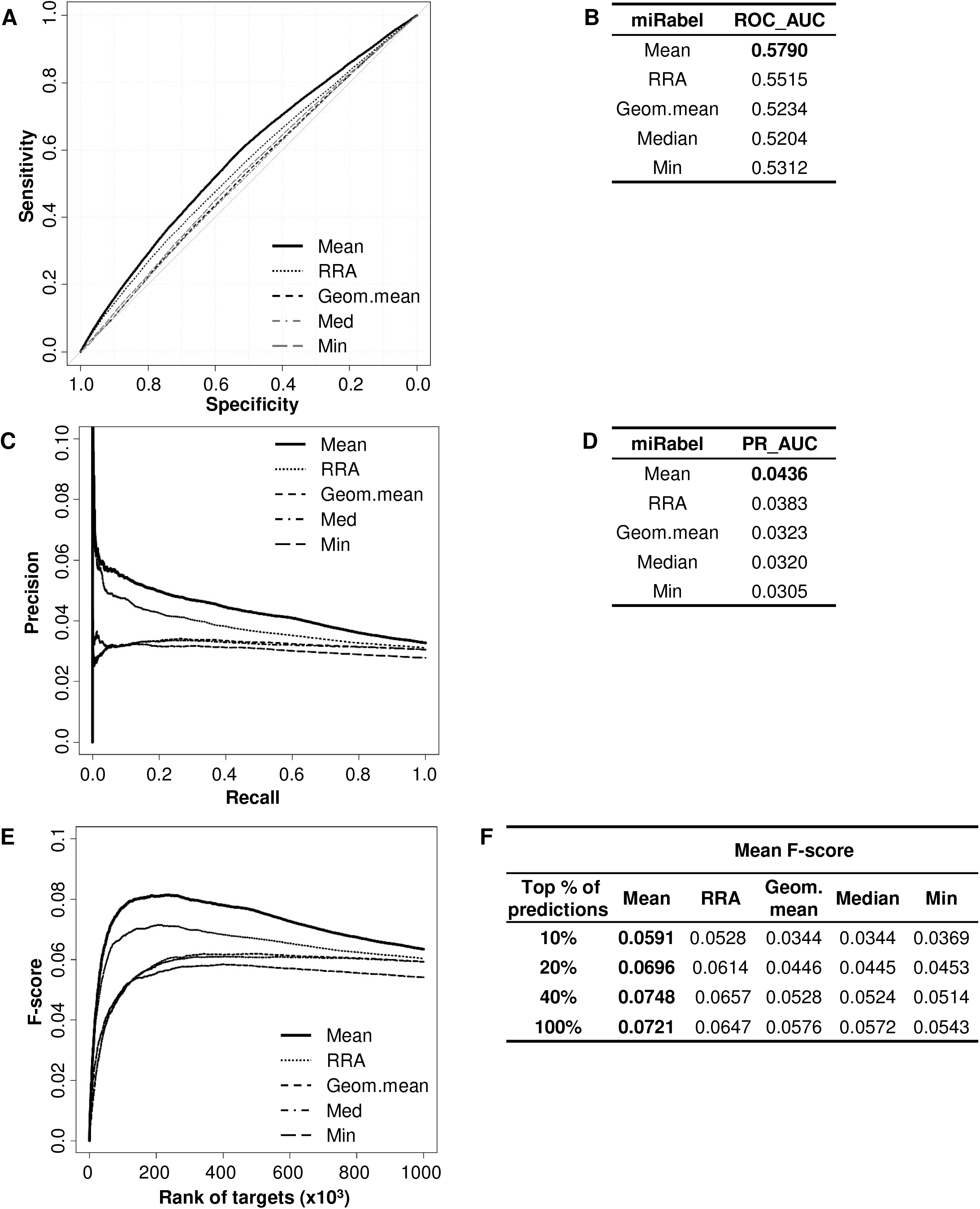
Performances comparison of aggregation methods. ROC curve analysis (**A**), showing the sensitivity and the specificity for 5 aggregation methods from the RRA R package, and their respective AUC (**B**) have been calculated using the pROC R package on 1 million random interactions. Using the same dataset, a precision and recall (PR) analysis (**C**) with PR_AUC (**D**) has been carried out using R programming as well. The cumulative harmonic mean between precision and recall (F-score) was also plotted (**E**) for each ranked interaction of this dataset. The average F-score is reported for the top 10%, 20%, 40% and all interactions (**F**). The higher are the ROC_AUC, PR_AUC and F-score, the better are the performances of the tested method. Highest values are in bold font.

### Comparison to aggregated methods

In order to test whether any improvement was gained with our aggregation method, the performances of each aggregated algorithms were compared to miRabel using ROC and PR analysis as well. These comparisons were done with 982,411 predicted interactions that are common to miRanda, PITA and SVMicrO. Within these predictions, 30,698 are experimentally validated ones. ROC curve analysis shows that miRabel improves the prediction of validated miRNA::mRNA interactions (ROC_AUC = 0.5984) compared to miRanda, PITA and SVMicrO (Fig. 4A-B). This improvement is even more visuable with the PR analysis (PR_AUC = 0.0437) (Fig. 4C-D) and the consistency of miRabel superior F-score throughout the dataset (Fig. 4E-F). Using 10 smaller datasets allowed us to confirm and to enhance the significativity of these analyses (p-value <10^−4^) (Table S3). A significant improvement was also manifest for the aggregated predictions for the top ranked interactions (ROC_pAUC90% = 0.0088; Sen90% = 0.1670) compared to miRanda, PITA and SVMicrO (Table S3).

**Figure 4:**
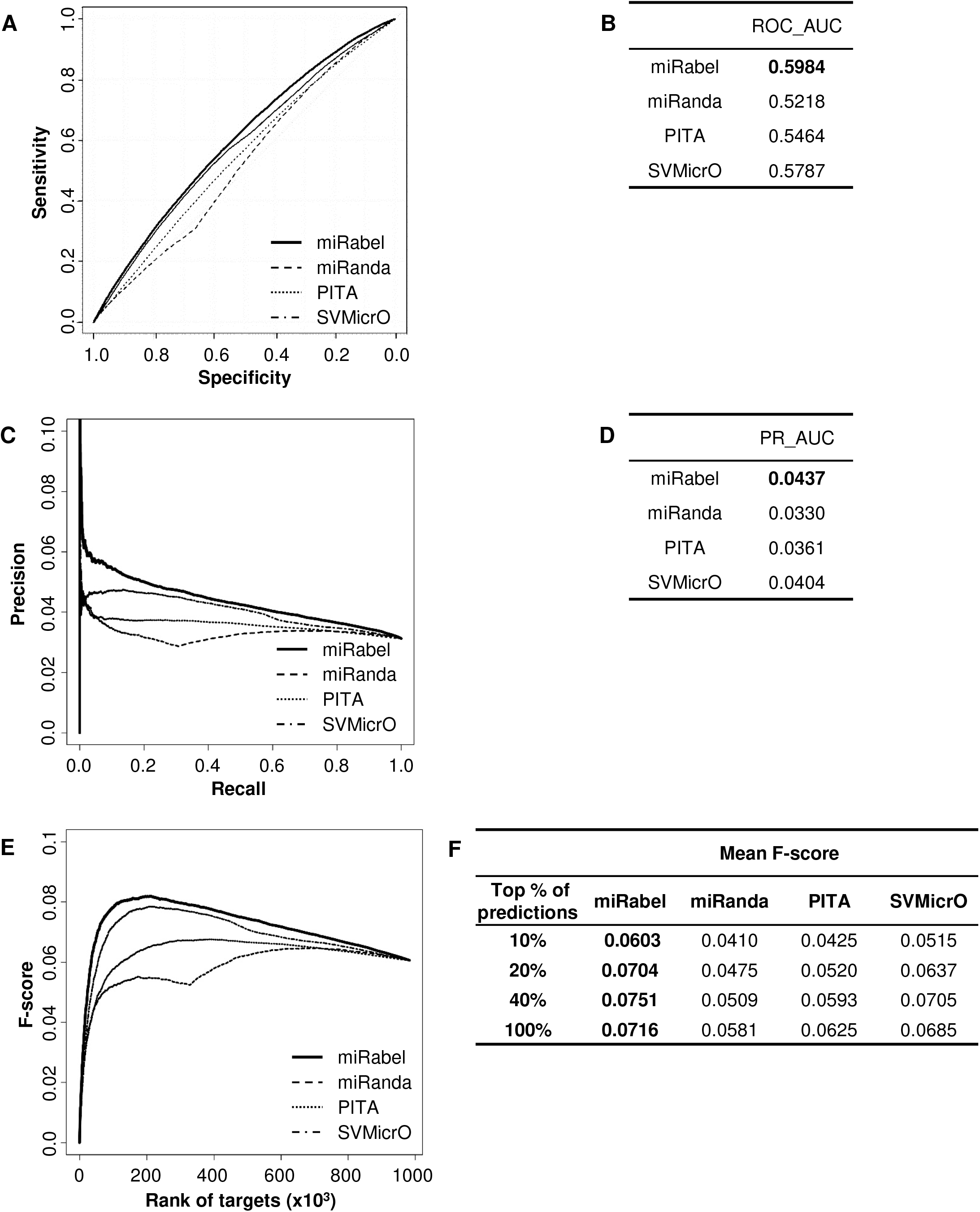
Performances comparison of aggregated prediction algorithms. ROC curve analysis (**A**), showing the sensitivity and the specificity for miRabel, miRanda, PITA and SVMicrO, and their respective AUC (**B**) have been calculated using the pROC R package on 982,411 common interactions. Using the same dataset, a precision and recall (PR) analysis (**C**) with PR_AUC (**D**) has been carried out using R programming as well. The cumulative harmonic mean between precision and recall (F-score) was also plotted (**E**) for each ranked interaction of this dataset. The average F-score is reported for the top 10%, 20%, 40% and all interactions (**F**). The higher are the ROC_AUC, PR_AUC and F-score, the better are the performances of the tested algorithm. Highest values are in bold font.

### Comparison to other prediction tools

The performances of miRabel were also compared to MBSTAR [37], miRWalk (v.2.0) [24], and TargetScan (v.7.1) [38], three efficient, up-to-date and/or widely used prediction web tools [21]. ROC and PR curves analysis using the same methods (all common interactions and ten random sets of 50,000 interactions) shows that our prediction data significantly improves the overall prediction of miRNAs target mRNAs when compared to MBSTAR (5 and Table S4) and miRWalk (6 and Table S5). However, even though miRabel shows better overall performance than Targetscan (ROC_AUC: 0.5577 vs 0.5477, p=3.5×10^−3^, Fig. 7-B, Table S6), they both seem fairly equal when we focus the analysis on true positives identification (PR_AUC: 0.0404 vs 0. 0406, Fig. 6C-F). Optimal specificity, ROC_pAUC90% and the corresponding sensitivity of our aggregated data exhibit also better performances than those of MBSTAR (Table S4) and miRWalk (Table S5) whereas these parameters are almost similar to the ones calculated for Targetscan (Table S6).

**Figure 5:**
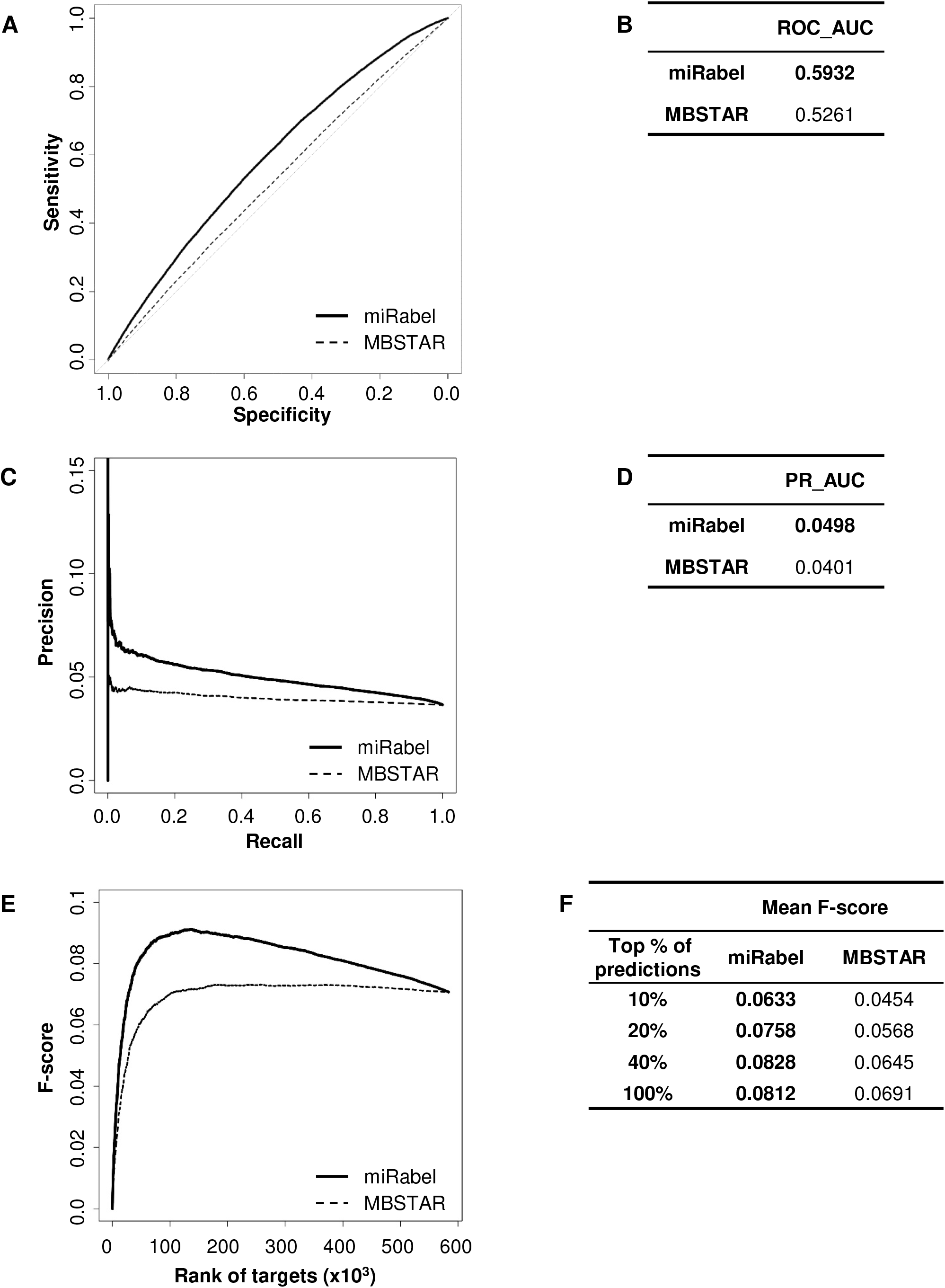
Performances comparison of miRabel and MBSTAR. ROC curve analysis (**A**), showing the sensitivity and the specificity for miRabel and MBSTAR, and their respective AUC (**B**) have been calculated using the pROC R package on 583,547 common interactions. Using the same dataset, a precision and recall (PR) analysis (**C**) with PR_AUC (**D**) has been carried out using R programming as well. The cumulative harmonic mean between precision and recall (F-score) was also plotted (**E**) for each ranked interaction of this dataset. The average F-score is reported the top 10%, 20%, 40% and all interactions (**F**). The higher are the ROC_AUC, PR_AUC and F-score, the better are the performances of the tested algorithm. Highest values are in bold font.

**Figure 6:**
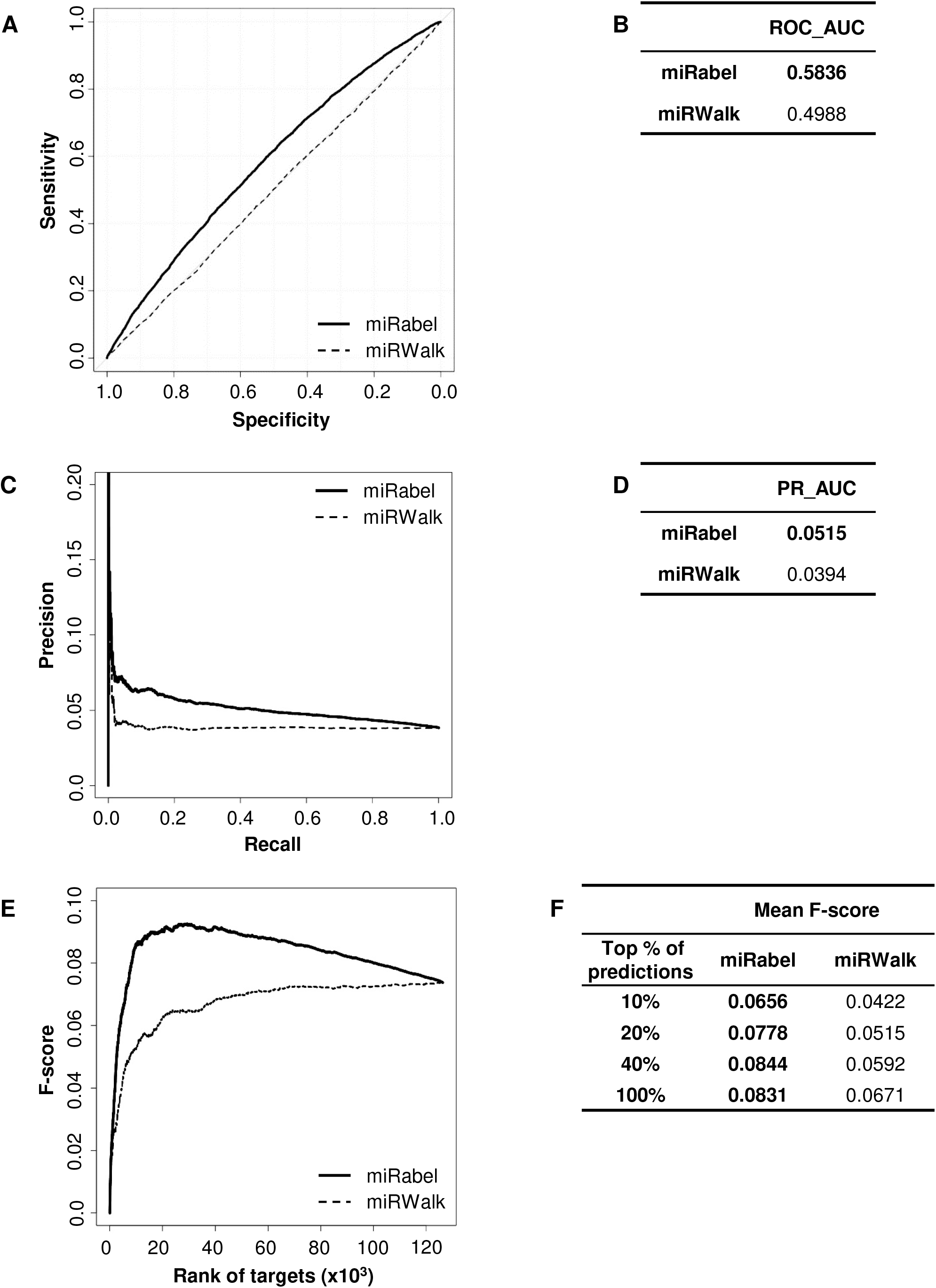
Performances comparison of miRabel and miRWalk. ROC curve analysis (**A**), showing the sensitivity and the specificity for miRabel and miRWalk, and their respective AUC (**B**) have been calculated using the pROC R package on 126,214 common interactions. Using the same dataset, a precision and recall (PR) analysis (**C**) with PR_AUC (**D**) has been carried out using R programming as well. The cumulative harmonic mean between precision and recall (F-score) was also plotted (**E**) for each ranked interaction of this dataset. The average F-score is reported the top 10%, 20%, 40% and all interactions (**F**). The higher are the ROC_AUC, PR_AUC and F-score, the better are the performances of the tested algorithm. Highest values are in bold font.

**Figure 7:**
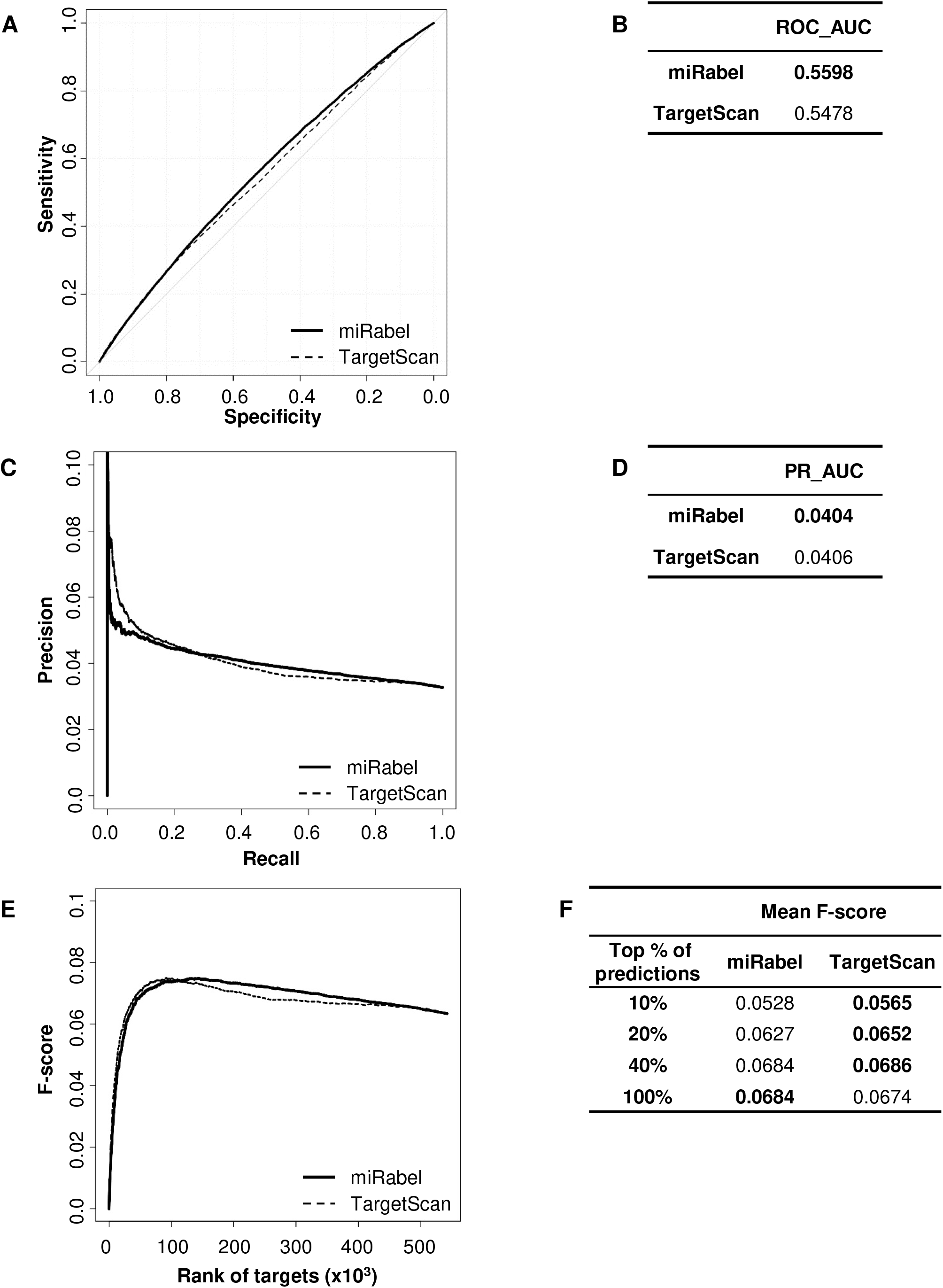
Performances comparison of miRabel and TargetScan. ROC curve analysis (**A**), showing the sensitivity and the specificity for miRabel and TargetScan, and their respective AUC (**B**) have been calculated using the pROC R package on 126,214 common interactions. Using the same dataset, a precision and recall (PR) analysis (**C**) with PR_AUC (**D**) has been carried out using R programming as well. The cumulative harmonic mean between precision and recall (F-score) was also plotted (**E**) for each ranked interaction of this dataset. The average F-score is reported the top 10%, 20%, 40% and all interactions (**F**). The higher are the ROC_AUC, PR_AUC and F-score, the better are the performances of the tested algorithm. Highest values are in bold font.

## Discussion

The prediction of miRNA targets is a bioinformatic challenge. Indeed, increased biological knowledge of the interactions between miRNAs and their targets has improved the predictions but they still suffer from high false positive rate [28]. Actually, each algorithm incorporates its own characteristics [39] and the comparison of their results highlights important contradictions in their respective predictions [39, 40]. We therefore hypothesized that the aggregation of the predictions of several algorithms would improve the relevance and the robustness of the prediction of miRNA targets.

In order to validate this concept, we have chosen to aggregate the predictions of three algorithms, miRanda, PITA and SVMicrO, because they use different but complementary information such as site accessibility or free energy to make their predictions. The results they provide are different both in terms of their probability of interaction (i.e., their ranking) and their number of target mRNAs [39]. Thus, only 11.4% of total interactions (982,411 / 8.6 million) are common to each other. The example of hsa-miR-16 that we present (Fig. 2B) also illustrates very well these divergences of predictions. Moreover, because these algorithms have not been updated recently, some more refined features of the seed region found in recent prediction approaches such as TarPmiR [41], are not considered in our aggregated results. This also explains why only 1,107 miRNAs have target mRNAs among the 2,578 that miRabel includes. Only the human miRNAs were used initially to limit the amount of data to be manipulated as well as the associated computation times, but the approach that we propose is generalizable to the miRNAs of all origins. Since the score generated by the RRA package is also representative of the significativity of the ranking for a given interaction, we suggest to use miRabel with a threshold of 0.05. Moreover, this is in agreement with the threshold estimated on the different ROC analyses using the closest top-left method (data not shown). We, however, acknowledge that further analyses are required to really define an optimal threshold for miRabel. Finally, the choice of algorithms is also limited by the free availability of their prediction database. To further improve predictions, it would therefore be interesting to take into account newer promising tools such as ComiR [42] or miRmap [43] whose prediction algorithms have been shown to perform well [39].

Comparing five of the aggregation methods included in the RRA package shows that the “mean” method is best for aggregating miRNA prediction lists (3, Table S2). However, although statistically significant, these values are relatively close to one another. These results are similar to those obtained in studies designed to compare the performance of several rank aggregation methods and showing better performances for the mean method [44-46]. Although not the best in our study, the RRA method can handle incomplete rankings and is robust to noise due to divergent lists [33]. In addition, it has already been used to aggregate miRNA profiles in a metaanalysis in nasopharyngeal cancer but without comparing it with other aggregation methods [47]. Among other aggregation methods, Cross Entropy Monte-Carlo has been found to be inadequate for our study due to too extensive computation times with large lists of items as previously reported [48]. As an example, a preliminary test showed us that it takes around 15 hours on a desktop computer for the ECMC method as integrated in the RankAggreg R package [49] to aggregate three short lists of only one hundred predicted mRNA targets from one microRNA (data not shown). Another method that could be evaluated with our data is the Borda count algorithm [50] which has already been used to aggregate cancer expression microarrays and proteomics datasets into a single optimal list [51].

Our miRNA target predictions database, called miRabel, performs better than each of the individual aggregated algorithms (4). Interestingly, prediction improvement is clearly visible in the top ranked interactions of miRabel (Table S3), thus showing that aggregating results from other tools moved validated interactions up in ranking and moved down less relevant ones. This is in line with multiple studies which show that combining data is so far the best compromise to obtain the most relevant interactions [16, 22, 40, 52, 53]. A recent study in particular shows that the union (but not the intersection) of the predictions of three tools among four (TargetScan, miRanda-mirSVR, RNA22) increases the performance of the analyses [54]. However, our work goes further since prediction lists were aggregated and re-ranked in a unique list. The performance of their method was evaluated using only ten miRNAs and 1,400 genes but not the entire database. In order to avoid selection bias of the datasets, we analyzed all 982,411 interactions common to miRabel and the three aggregated algorithms, which represent 519 miRNAs and 14,319 genes. The use of ten random datasets of 50,000 interactions also enhances the relevance and statistical analysis of the results. Furthermore, even though miRabel aggregates older databases, it shows equal (vs. TargetScan) or better (vs. MBSTAR and miRWalk) performances than up-to-date algorithms, thus clearly establishing that our method, even though simple, has a great potential. Interestingly, from all evaluations done with our datasets and methodology, we found that other algorithm performances to be quite different from what was originally described in their respective original publications. This is in agreement with a previous study which highlighted the importance of testing prediction results on multiple, independent datasets and with a standardized evaluation protocol [39]. This is also one of the strengths of our study. Indeed, throughout all comparisons, miRabel was tested on 55 different datasets, which gives more robustness to the performance values calculated for our method.

## Conclusions

MiRabel is a new efficient tool for the prediction of miRNA target mRNAs and their associated biological functions. Using an aggregation method, we improved the relevance of the predictions of 3 available algorithms. This promising approach can easily be extended to all publicly available databases or to other species. Moreover, the integrated biological pathways provide a more comprehensive view and new insights into the complex regulatory network of miRNAs.

## Acknowledgements

We thank Dr. Isabelle Lihrmann for critical reading of the manuscript and fruitful discussion. We are grateful to the CRIANN (Centre Régional Informatique et d’Applications Numériques de Normandie) for allowing us to use their computing facility and University of Rouen Normandy to host the miRabel’s website.

